# The most efficient metazoan swimmer creates a ‘virtual wall’ to enhance performance

**DOI:** 10.1101/2020.05.01.069518

**Authors:** Brad J. Gemmell, Kevin T. Du Clos, Sean P. Colin, Kelly R. Sutherland, John H. Costello

## Abstract

It has been well documented that animals (and machines) swimming or flying near a solid boundary get a boost in performance^1-6^. This ground effect is often modeled as an interaction between a mirrored pair of vortices represented by a true vortex and an opposite sign ‘virtual vortex’ on the other side of the wall^7^. However, most animals do not swim near solid surfaces and thus near body vortex-vortex interactions in open-water swimmers have been poorly investigated. In this study we examine the most energetically efficient metazoan swimmer known to date, the jellyfish *Aurelia aurita*, to elucidate the role that vortex interactions can play in animals that swim away from solid boundaries. We used high speed video tracking, laser-based digital particle image velocimetry (dPIV) and an algorithm for extracting pressure fields from flow velocity vectors to quantify swimming performance and the effect of near body vortex-vortex interactions. Here we show that a vortex ring (stopping vortex), created underneath the animal during the previous swim cycle, is critical for increasing propulsive performance. This well positioned stopping vortex acts in the same way as a virtual vortex during wall-effect performance enhancement, by helping converge fluid at the underside of the propulsive surface and generating significantly higher pressures which result in greater thrust. These findings advocate that jellyfish can generate a wall-effect boost in open water by creating what amounts to a ‘virtual wall’ between two real, opposite sign vortex rings. This explains the significant propulsive advantage jellyfish possess over other metazoans and represents important implications for bio-engineered propulsion systems.

---

The phenomenon of increased lift generated over static surfaces moving parallel to solid boundary is termed ‘steady ground’ or ‘steady wall’ effects. This process has been widely studied and reviewed^8^ leading to a broad literature describing the advantages of gliding both in air^1-4^ and water^5,6^. The increase in lift near the ground is largely attributed to decelerated flow beneath the lifting surface resulting in higher underside pressures. This effect is often modeled as an interaction between a mirrored pair of vortices represented by a true vortex and an opposite sign ‘virtual vortex’ on the other side of the wall^7^. More recently, it has been shown that considerable performance benefits also exist for non-static, active swimming near a solid boundary. For example swimming animals exhibit reduced cost of transport and improved energetic efficiencies when near a wall^9,10^. Fernández-Prats, et al. ^11^ found considerable propulsive advantages for an undulatory, near-wall swimmer with a 25% increase in speed and 45% increase in thrust near a solid surface. Thus the increase in lift for a static propulsor translates to an increase in thrust in the unsteady case^12,13^. However, this phenomenon has not been explored for unsteady propulsors away from solid boundaries even though numerical models have suggested true vortex-vortex interactions should produce an equivalent effect.

Jellyfish represent an intriguing target for investigations of open water vortex-vortex interactions as they are the most energetically efficient swimmers known to date^14,15^ and have variety of traits that make them ideal organisms for fluid dynamics work (see supplemental information for more details). These animals can utilize passive energy recapture (PER) of a single vortex ring (stopping vortex) allowing the moon jellyfish (*Aurelia aurita*) to travel 30% further per swim cycle^15^. In this study we explore the role that vortex-vortex interactions play in medusan swimming. We demonstrate a previously undescribed role of the stopping vortex with respect to its ability to enhance thrust and swimming performance as it interacts another vortex ring to generate a ‘virtual wall effect’. The result is a significant increase in overall swimming performance and may also have important implications for understanding feeding in medusae, locomotion in other taxonomic groups as well as informing design principles for the engineering of bio-inspired vehicles.

Swimming kinematics differ greatly in derived taxa such as fish depending on whether they starting from rest or are at steady state^16-18^. Conversely, medusae display no difference in kinematics (Fig 1, Suppl. Table 1). This is due to the inner nerve ring network of neurons producing all-or-none overshooting action potentials that precede each swimming contraction ^19^. This generation of consistent swimming kinematics whether starting from rest or already undergoing steady swimming provides a unique opportunity to explore the role of vortex interactions in biological propulsion. While kinematics did not differ, we observed significant performance differences in the swim cycles between animals beginning from rest and those already swimming (Fig 1b, Suppl. Table 1). Animals that were already swimming saw a 41% increase in maximum swimming speed and an 61% increase in cumulative distance travelled per swimming cycle compared to those starting from rest. This performance discrepancy is unlikely due to any differences in momentum between the two types of swimming cases as small *A. aurita* come to a stop (near zero forward velocity) at the end of each swimming cycle (Figure 1b). This period lasts for approximately 0.5 s. Thus, whether starting from rest or during steady swimming, small *A. aurita* begin each new swim cycle with negligible forward momentum.

**Figure 1.**
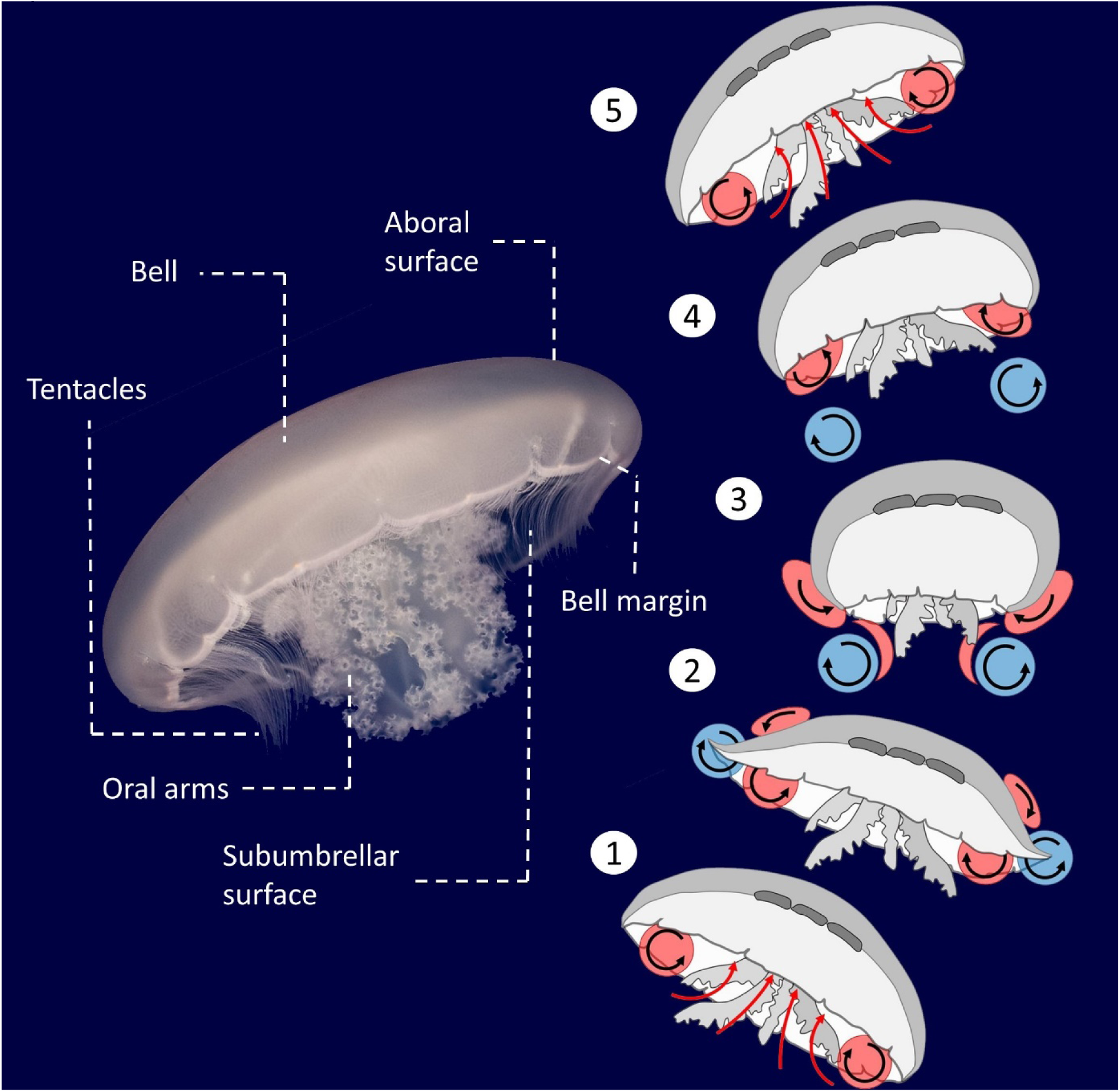
Anatomical features of a jellyfish and the vortex arrangement over the course of a swim cycle for the moon jellyfish (*Aurelia aurita*). Red = stopping vortex, blue = starting vortex. Black arrows = direction of fluid circulation. 1 and 5) Passive energy recapture mechanism (Gemmell et al. 2013). 2) Utilization of suction thrust from the upstream stopping vortex (Colin et al. 2012; Gemmell et al. 2015). The significance of the early stages of vortex-vortex interactions at the subumbrellar surface has not been investigated and is the focus of the present study. 3) Completion of the bell contraction. The starting vortex travels downstream in the wake after cancellation of the stopping vortex (Dabiri et al. 2007). 4) The stopping vortex is enhanced and repositioned inside the subumbrellar cavity (Gemmell et al. 2014). The newly formed stopping vortex and the downstream starting vortex interact to form a laterally oriented vortex superstructure (Dabiri et al. 2005).

**Figure 1.**
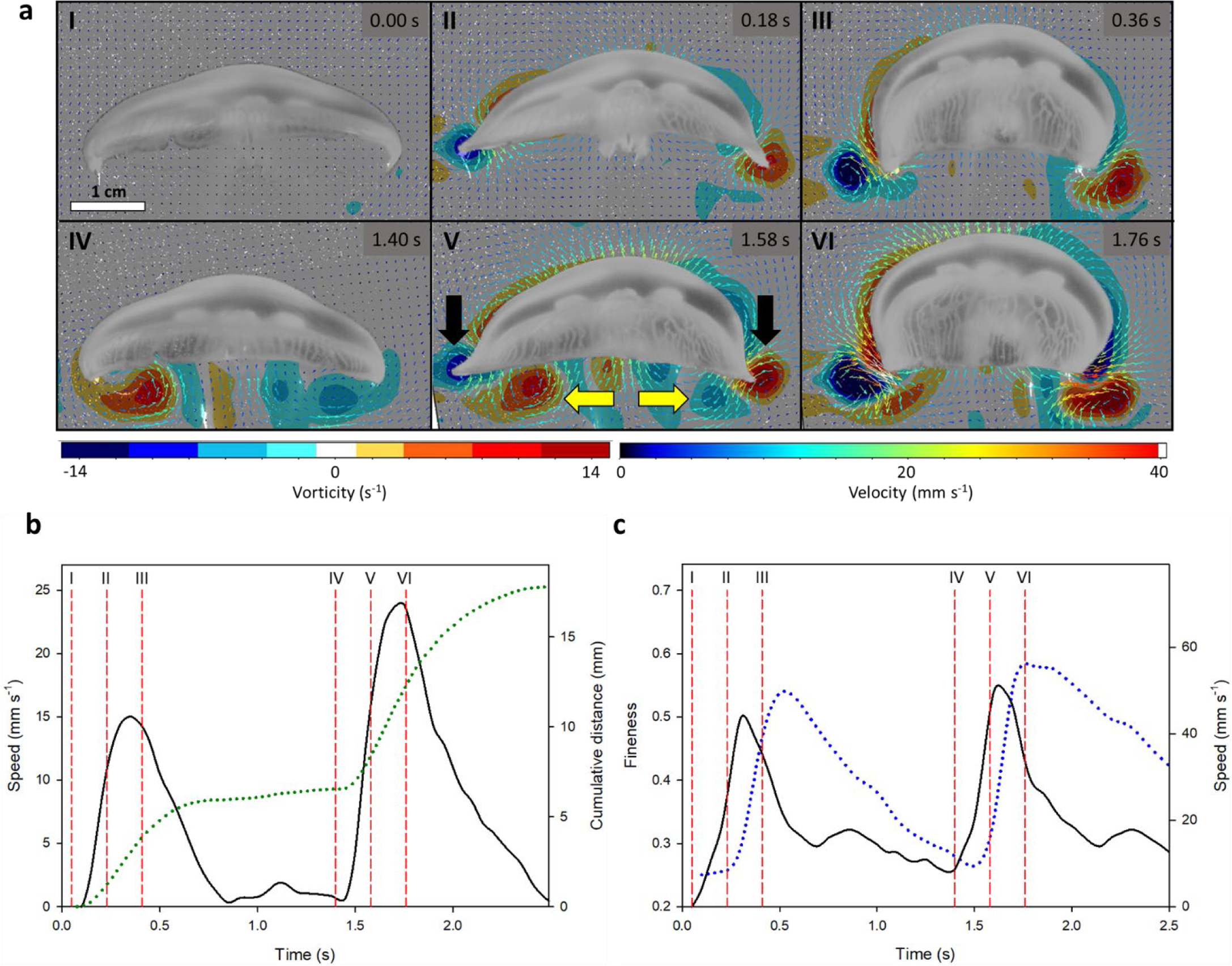
Representative swimming sequence of a 4 cm moon jellyfish (*Aurelia aurita*) as it begins swimming from rest. a) Fluid vorticity and velocity variables. Note the lack of vorticity under the bell in panel I and the presence of a stopping vortex prior the next second contraction cycle in panel IV. Black arrows show an example of the starting vortex and yellow arrows show the stopping vortex. b-c) Kinematic and performance variables during two swim cycles shown in panel (a). Vertical dashed lines (red) denote the panel numbers in Figure (a). b) Swimming speed (solid black line) and cumulative distance (dotted line). c) Bell fineness (blue dotted line) and instantaneous speed of the bell margin (solid line).

Therefore, in jellyfish, the only notable difference between the two swimming cases is the presence or absence of a stopping vortex underneath the bell of the medusa. Individuals starting from rest display no vortices while large stopping vortices (mean maximum vorticity: 18 s^−1^ s.d. 4) were evident for individuals that had just completed a swimming cycle (Figure 1). This stopping vortex persists into the beginning of the next swim cycle maximum vorticity values of 11 s^−1^ (s.d. 3) are observed. Vorticity in this same region for animals starting from rest was near zero (0.2 s^−1^ s.d. 0.1).

As a swim cycle is initiated, a vortex ring termed a starting vortex forms at the tip of the bell margin (Fig 1) and has an opposite sign to the stopping vortex. In steady swimming cases the bell margin and resulting starting vortex are forced towards and begin to interact with the opposite sign stopping vortex. This creates a zone of convergent flow directly underneath the bell margin (Fig 2). After the contraction phase the jellyfish’s bell resets during the relaxation phase of the swim cycle. The upward and outward movement of the bell margin both enhances the strength of the stopping vortex and keeps it positioned close to the subumbrellar surface^20^. The rotation of the stopping vortex as it sits under the animal drives fluid against the subumbrellar surface which contributes to the passive energy recapture effect^15^. In *A. aurita*, the next contraction cycle begins before the stopping vortex dissipates (Figures 1, 2). This overlap in space and time of the starting and stopping vortices results in the two vortices interacting at the ventral side of the propulsor. Fluid is driven towards the subumbrellar surface from the stopping vortex and away from the subumbrellar surface by the movement of the bell margin itself (Figure 2, 4b).

**Figure 2.**
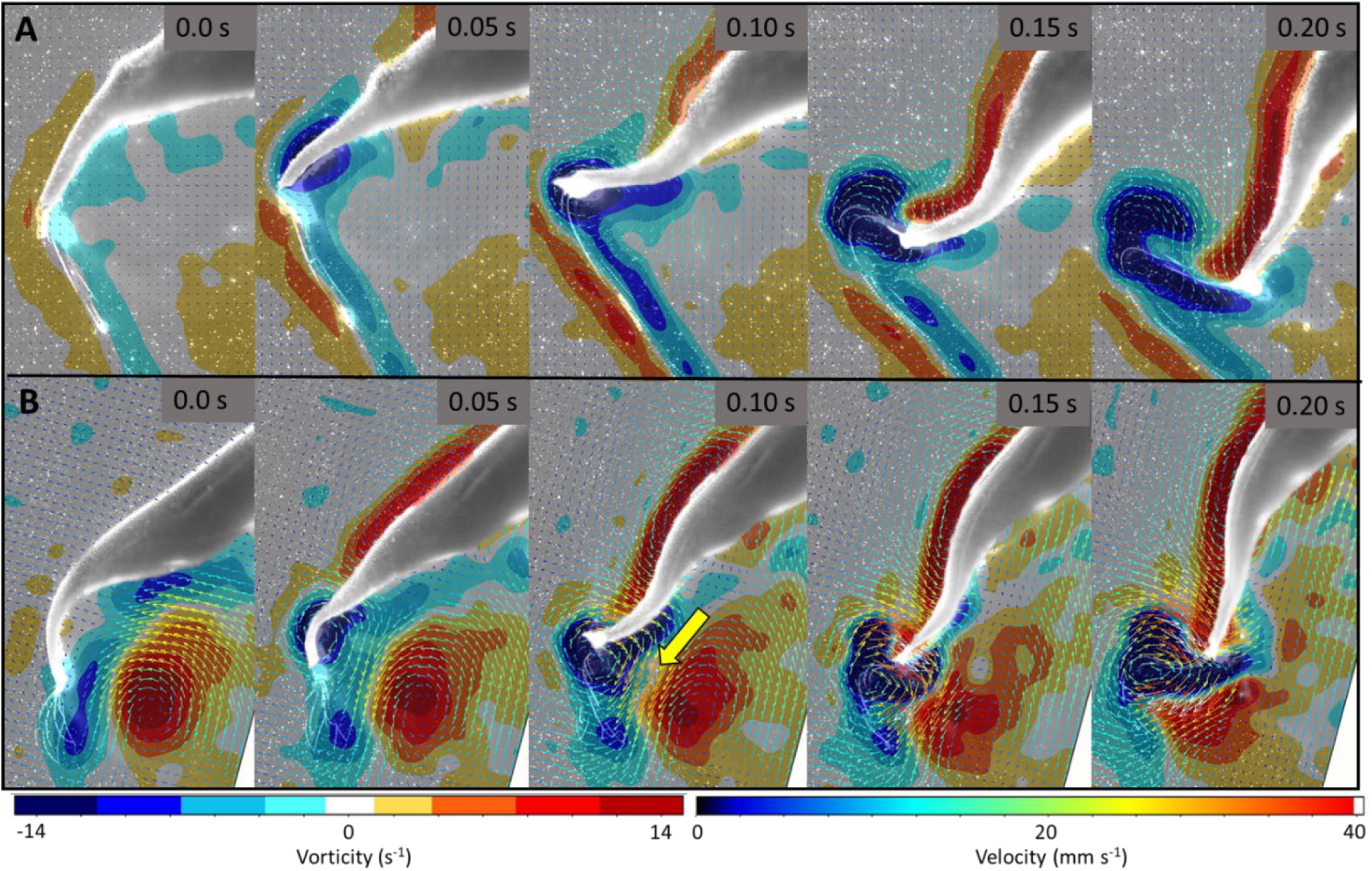
Fluid vorticity and velocity vectors around the bell margin for a representative jellyfish swimming sequence when the animal begins from rest (A) and when undergoing steady swimming (B). Yellow arrow indicates region of highest flow due to vortex interface acceleration (VIA). This only occurs when a stopping vortex is present under the animal.

**Figure 3.**
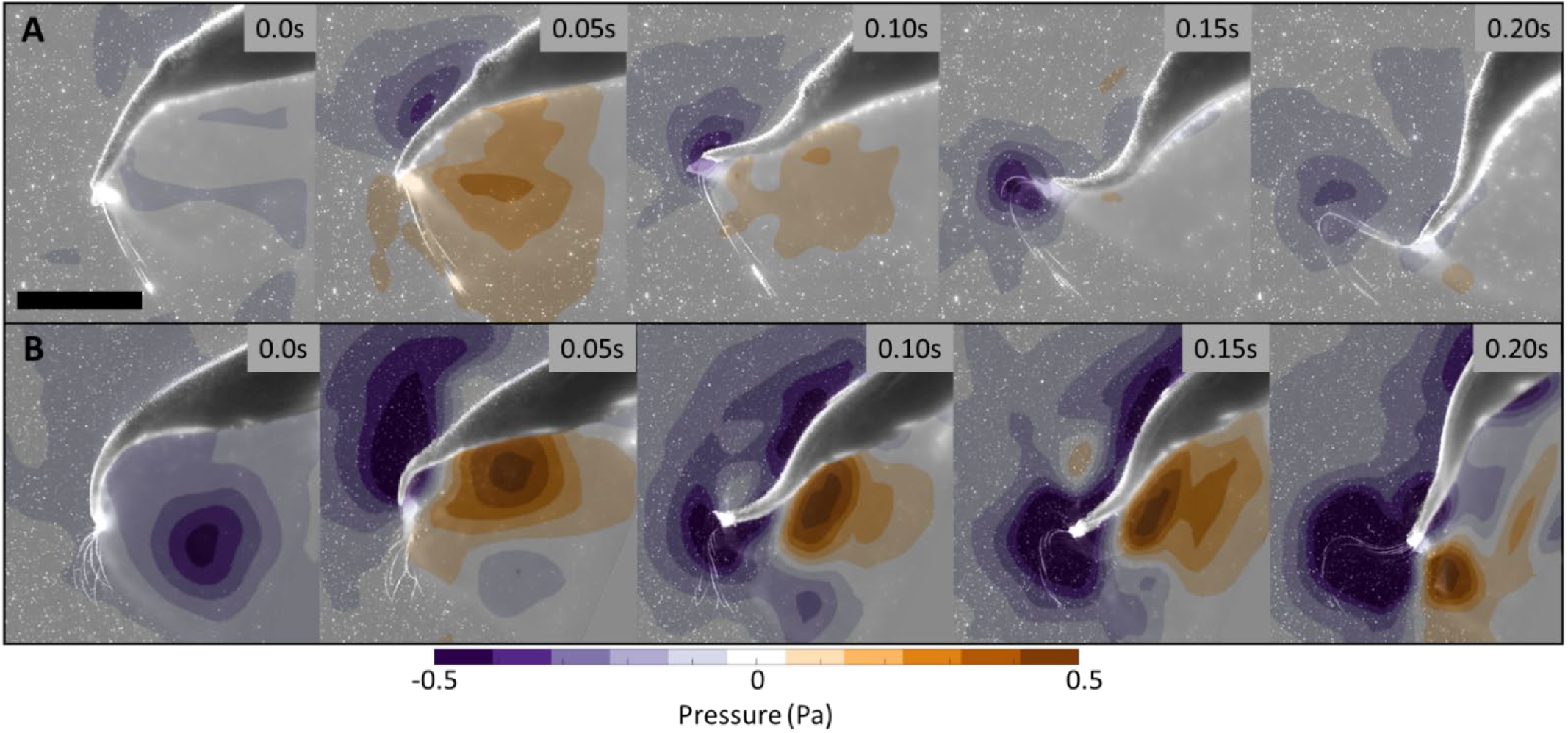
Pressure fields calculated from the same jellyfish swimming sequences as in Figure 3 when the animal begins from rest (A) and when undergoing steady swimming (B). Scale bar = 0.5 cm.

**Figure 4.**
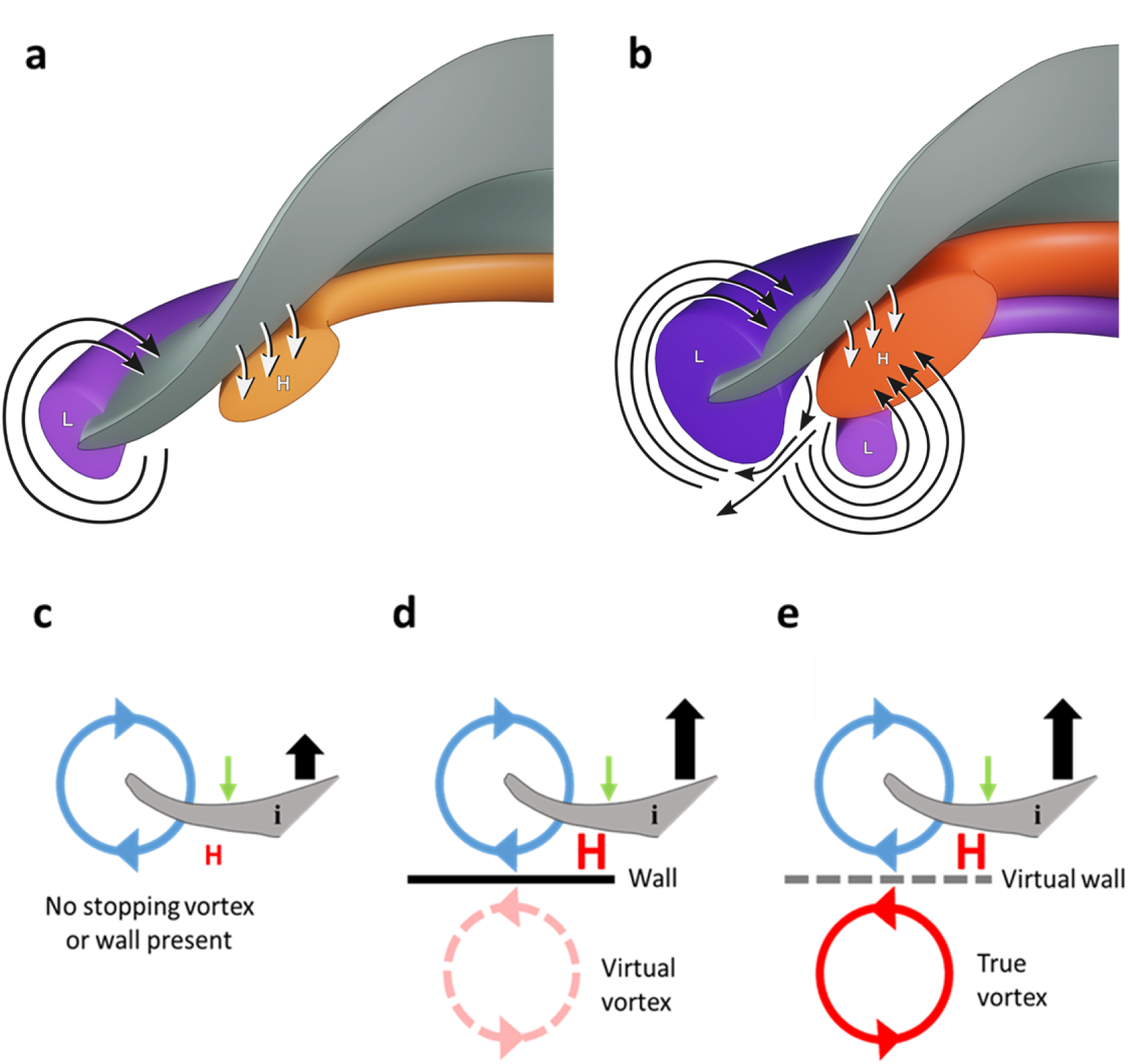
Diagrammatic representations of water movement and pressure anomalies due to kinematic movement of the jellyfish bell. Note: a cross section through only one half of the animal is depicted (grey surface) for simplicity. a) Jellyfish swimming without a developed stopping vortex underneath the bell. b) Jellyfish swimming with a developed stopping vortex underneath the bell. Red areas denote regions of above ambient pressure and blue regions denote regions of below ambient pressure. Darker colors represent higher magnitudes of pressure. Blue lines show the direction of fluid motion. Green arrows show the direction of movement of the bell margin during the contraction phase and black arrows depict the relative forward thrust generated. c) Away from any solid boundary and without the presence of an additional opposite sign vortex. d) Ground effect case where swimming near a solid boundary produces an opposite sign virtual vortex and results in greater positive pressures compared to panel A. Adapted from ^7^. e) Away from a solid boundary with a real opposite sign stopping vortex present. This case represents a jellyfish undergoing routine swimming and results in a “virtual ground effect” thrust benefit similar to panel B. Jellyfish bell margin represented by symbol (i). Green arrows denote direction of propulsor motion and black arrows denote relative thrust generation.

In order to determine the role that vortex-vortex interactions plays in jellyfish swimming, we calculated the pressure fields around the bell margin of swimming medusae. In cases where the stopping vortex was absent, pressure fields near the bell margin were near ambient just prior to contraction (Figure 3). Conversely, swimming cases where the stopping vortex was present prior to bell contraction saw a region of negative pressure at the location of the stopping vortex (Figure 3). The mean peak negative pressure in this region just prior to bell contraction was −0.35 Pa (n=5, s.d. 0.17). Within 100 ms after bell contraction is initiated, a strong positive pressure region appeared at the subumbrellar surface of the bell margin (Figure 3). Peak positive pressures in this region reached 0.74 Pa (n=5, s.d. 0.08). These positive pressures were significantly stronger (T-test, P < 0.001, n=5) when a stopping vortex was present compared to swim cycles without a stopping vortex (0.32 Pa, n=5, s.d. 0.09). Peak negative pressure values were not significantly different (T-test, P = 0.120, n=5).

The magnitude of the pressure gradient on either side of a propulsor is directly related to the thrust generated^21-23^. Thus, the greater positive pressures on the underside of the bell margin when a stopping vortex is present provides medusae with additional thrust without the need to alter swimming kinematics. In is interesting to note that our observations of swimming speed enhancement of 41% when a vortex-vortex interaction occurs near the propulsor tip of jellyfish is nearly identical to the maximum 40% thrust enhancement observed during ground effect experiments of a pitching foil near a solid wall^13^.

The similarities between performance enhancement in jellyfish when vortex-vortex interactions occur and that of near-wall swimmers is striking considering jellyfish are achieving these benefits well away from any solid surface. This can be resolved by the fact that the ground effect is realized with a simple symmetry approach by computing the flow around the object and its virtual mirror image, considering the symmetry-line as the ground^7^. In the case of jellyfish, the animals simply appear to be substituting the mirrored virtual vortex with a real one, thus making the ground or wall virtual instead (Fig 4). We thus describe the jellyfish circumstance as a ‘virtual wall’ effect. Engineers frequency consider ways to employ ground effect^24,25^ because it can greatly improve thrust and efficiency of a propulsor, however the requirement of propelling very near a solid boundary limits potential applications. The ability to arrange vortices to create a ‘virtual wall’ as in jellyfish swimming, demonstrates a currently unrecognized potential for future engineered vehicles to benefit from a ground effect without actually requiring the ground.

The development of jellyfish as a model to understand effective propulsion though a fluid has many potential benefits (see supplemental discussion for details). Unlike other swimmers such as crustaceans, cephalopods and fish, jellyfish are constrained by having muscle tissue that is only a single cell mono-layer thick^26^. This limits the power that medusae can create when swimming. However, given this resource limitation of power output available for propulsion, there could have been enhanced selective pressure^27^ in this group of animals to develop and refine mechanisms to organize fluid and vortices for maximal performance. It is currently unknown how widespread vortex-vortex interactions with animal propulsive structures are in nature, as we are aware of very few other examples^28^. However, given observations of wake capture in insects^29^ and the substantial interest in developing bio-inspired vehicles, especially those based on a jellyfish platform^30-32^, this represents an important direction for future investigation.

## Methods

Jellyfish (*Aurelia aurita*) were obtained from the New England Aquarium and maintained in the laboratory in 20L aquaria and fed with newly hatched *Artemia salina* nauplii daily. Free-swimming jellyfish between 3.8 and 4.2 cm in diameter (n=8) were recorded in a glass filming vessel (30 × 10 × 25 cm) by a high-speed digital video camera (Fastcam 1024 PCI; Photron) at 1,000 frames per second. Only recordings of animals swimming upward were used in the analysis to eliminate the possibility of gravitational force aiding forward motion of the animal between pulses. Detailed kinematics (2D) were obtained using Image J v1.46 software (National Institutes of Health) to track the *x* and *y* coordinates of the apex of the jellyfish bell and the tips of the bell margin over time. Swimming speed was calculated from the change in the position of the apex over time as:

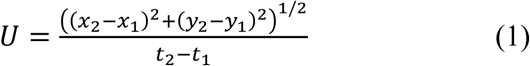

Jellyfish were illuminated with a laser sheet (680 nm, 2W continuous wave; LaVision) oriented perpendicular to the camera’s optical axis to provide a distinctive body outline for image analysis and to ensure the animal remained in-plane, which ensures accuracy of 2D estimates of position and velocity. Kinematic data were log-transformed and checked for normality using a Shapiro– Wilks test. Data were subsequently tested using one-way ANOVA to determine if a significant difference existed between means.

Fluid motion created by the jellyfish while swimming was quantified using 2D digital particle image velocimetry. Using the setup described above, the filtered seawater was seeded with 10 μm hollow glass beads (Dantec Dynamics). The velocities of particles illuminated in the laser sheet were determined from sequential images analyzed using a cross-correlation algorithm (LaVision software). Image pairs were analyzed with shifting overlapping interrogation windows of a decreasing size of 64 × 64 pixels to 32 × 32 pixels or 32 × 32 pixels to 16 × 16 pixels.

Pressure fields calculations were based on the queen 2.0 pressure field calculation package for MATLAB^33^. Pressure field data was inferred from the measured velocity fields by numerically integrating the inviscid Navier-Stokes equation, or Euler equation:

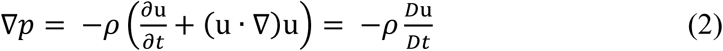

where ρ is the fluid density and **u** is the Eulerian velocity field. Details of this method are given in ^34^ and described briefly below.

The material acceleration term 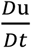, which quantifies the acceleration of individual fluid particles in the flow, was calculated from the measured DPIV velocity field **u**(*x, y*). The pressure term was then determined to within a constant of integration by integrating equation 2 spatially. To reduce errors in the numerical integration of the measured velocity data, the procedure of Liu and Katz ^35^ was employed. Data were input a time series of DPIV data on a 128×128 grid. Preprocessing of the DPIV data was completed in MATLAB to compute the material acceleration 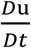, which is also a required input to the pressure calculation code. Material acceleration was determined by computing the difference in the velocity of fluid particles initially located at the DPIV data grid points at time *t1* and subsequently advected to new positions time *t2* ^34^. The output data from the code is a time series of pressure fields with scalar pressure computed at each of the 128×128 nodes of the corresponding DPIV fields.

## Acknowledgements

The authors thank John Dabiri and Megan Leftwich for insightful discussions regarding vortex dynamics. This research was supported by grants from the National Science Foundation (CBET-1511996 and OCE-1829945) to BJG and (OCE-1536688 and OCE-1829913) to SPC and JHC.

## Author contributions

BJG, SPC and JHC designed the experiments. BJG collected the data. BJG, KTD and KRS analyzed the data. BJG wrote the initial draft of the manuscript and all authors contributed to revising the manuscript.

## Data availability

The datasets generated during and/or analyzed during the current study are available from the corresponding author on reasonable request.

## Competing interests

Authors declare no competing interests.

## Notes

### Competing Interest Statement

The authors have declared no competing interest.

