## supplemental text and table for "The most efficient metazoan swimmer creates a ‘virtual wall’ to enhance performance"

**Supplementary Information**

Supplementary background information:

As a taxonomic group, jellyfish have been swimming the world's oceans for over 500 million years (*1*) and are the most energetically efficient swimmers known to date (*2, 3*). Their high performance relies on generating effective body-fluid interactions despite having muscle tissue that is only a single cell mono-layer in thickness (*4, 5*). The ability to generate sufficient thrust with very low power input makes medusae an intriguing target for elucidating the role of body kinematics and resulting fluid interactions. Other characteristics of medusan swimmers such as their axisymmetric morphology, translucent body tissues and consistent swimming kinematics make them an ideal animal model to investigate questions of body flexibility, vortex interactions and their roles in generating favorable pressure fields for thrust.

Animals are able to move effectively through fluids by transferring the momentum of body movements to the surrounding fluid in a manner that efficiently produces and controls thrust production. The generation and control of vortices along the body is critical for this effective movement though a fluid (*6-9*). An interesting characteristic that appears to be universal of swimming and flying animals is that their propulsors are flexible and, across taxa, these propulsors bend with strikingly similar kinematics (*10, 11*). This is in sharp contrast to traditional human-generated propulsors which have an almost universally rigid configuration. Bending is thus critical for animal thrust production and studies have shown that flexible propulsors increase thrust production up to an optimal level, beyond which further bending reduces thrust (*12-16*).

One effect of bending is the enhancement of negative pressure regions in the fluid adjacent to the inflexion point of the bending appendage (*13, 17, 18*). Generating a strong pressure gradient across an appendage is critical for creating substantial thrust. Stronger pressure fields can accelerate more fluid around propulsors, which enhances the momentum of the surrounding fluid. The pressure gradients that will generate the most thrust will have a substantial negative pressure on one side of the appendage and a strong positive pressure field on the other. However, the mechanistic explanations for the variability in appendage kinematics, optimization of pressure fields, thrust production and the fluid interactions that drive them are not well understood.

Earlier work on jellyfish locomotion recognized that vortex formation (i.e. starting and stopping vortices) plays an important role in rowing-based medusan propulsion (*4, 19-21*) but the mechanistic underpinnings of the vortex-vortex interactions and resulting impact on thrust generation were not well understood. However, more recent work demonstrated that jellyfish can utilize passive energy recapture (PER) of the stopping vortex (*3*) allowing the moon jellyfish (*Aurelia aurita*) to gain an additional 30% more distance per swim cycle without the input of additional energy. This PER is accomplished by the enhancement and repositioning of a stopping vortex during the refilling (i.e. recovery) phase of the swim cycle by the flexible margin of the medusa bell (*22*). The jellyfish then pauses before the next swimming contraction and the induced flow of the stopping vortex ring creates a high pressure region at the subumbreallar surface thereby contributing additional thrust to move the animal forward (*3*). The previously unrecognized importance of the stopping vortex is likely due to the fact that this vortex ring remains obscured within the medusan subumbrella until contraction of the jellyfish bell occurs and the remnants of stopping vortex are ejected with the starting vortex, which have merged into a laterally oriented vortex superstructure (*23*).

The use of the stopping vortex to gain additional distance each swim cycle is widespread in medusae (*24*). We have previously shown that while the concealed stopping vortex allows jellyfish to travel greater distances each swimming cycle, the overall benefit in terms of distance travelled primarily depends on the pause duration before the next cycle of bell contraction begins. It should be noted that across a diverse array of jellyfish taxa, the jellyfish begins the next swimming cycle well before the stopping vortex dissipates (*3, 24*). Thus, the maximum potential of the stopping vortex is not fully exploited and therefore limits the overall distance gained through the PER mechanism. It has remained unclear why this occurs and why jellyfish do not take full apparent advantage of the stopping vortex.

Results:

While the starting vortex is always present during *A. aurita* swimming, the strength in terms of vorticity of the starting vortices was significantly lower (T-test, $P = 0.02$, $n=8$) for animals starting from rest. This is despite having the same bell kinematics as steady swimmers with peak vorticity values of 18 s-1 (s.d. 3) and 25 s-1 (s.d. 5) respectively (Figure 2). The stronger starting vortices in steady swimmers coincide with the presence of opposite sign stopping vortices that were formed during the previous swimming cycle (Figure 2).

The highest fluid speeds were observed at the interface of these opposite sign vortices, at the subumbrellar surface of the bell (Figure 3) and particle tracking showed that at least some of the moving fluid that was accelerated between starting and stopping vortices ended up entrained into the growing starting vortex during contraction of the jellyfish bell. For cases where no stopping vortices were present prior to bell contraction, the highest fluid velocities were observed on the aboral side of the bell margin (Figure 3a) and the strong convergent flows at the subumbrellar surface were not observed.

Peak negative pressures around the bell margin of -0.34 Pa (n=5, s.d. 0.11) were observed when no stopping vortex was present and reached a peak value of -0.51 Pa (n=5, s.d. 0.12) when a stopping vortex was present. However, peak negative pressures were not significantly different (T-test, P = 0.120, n=5).

### Supplemental Discussion

Despite having translucent tissues, details of the stopping vortex are not easy to resolve in many species. Most scyphozoan medusae tissues are simply too opaque to accurately resolve flow tracer particles near the subumbrellar body surface. Only in smaller *Aurelia aurita* (> 6 cm) and only at the ideal body orientation were we able to resolve the entirety of the stopping vortex underneath the animal's bell with sufficient resolution for analysis. Even with the challenges of observing all of the near-body vortices during swimming, medusae provide an ideal biological platform for investigating the role of vortex interactions for effective propulsion. The ability to resolve flows around the entire body and axisymmetric morphology allow for 2-dimensational methods (i.e. laser-based dPIV) to provide accurate, detailed quantification of flows near the surface of propulsive structures. Furthermore, compared to more derived taxa such as fish, where swimming kinematics often differ greatly whether starting from rest or steadily swimming (*27-29*), medusae display a much more limited range of motion. This is due to the fact that the inner nerve ring network of neurons produce all-or-none overshooting action potentials that precede each swimming contraction (*30*), thereby limiting the range of kinematic variation during swimming. The generation of consistent swimming kinematics whether starting from rest or already undergoing steady swimming provides a unique opportunity to explore the role of vortex interactions in biological propulsion.

As the medusa contracts its bell, the starting vortex is forced closer and closer to the stopping vortex causing a strong jet to form at the interface and quickly reducing the magnitude of the stopping vortex as rotational energy is translated into the straight jet (Figure 2). This straight jet is known as the vortex interface acceleration (VIA) (*31*) and its formation appears to coincide with thrust in several types of swimmers (*31, 32*). At the interface of the starting and stopping vortices as rotational momentum is converted to a straight jet momentum and accelerates the overall flow adjacent to the bell margin. Through particle tracking we observed that some of the fluid from this high velocity jet is incorporated into the starting vortex as bell contraction progresses. Thus, by incorporating this high speed fluid into the starting vortex, it appears that medusae can generate stronger vortices and greater negative pressures without significantly altering the bell margin kinematics and presumably energetics.

An example of what happens as a result of forcing vortices of opposite sign together can be seen in the case of vortex ring approaching a solid wall. This highly replicable and well-studied phenomenon can provide some insights into effective biological propulsion. As a vortex ring approaches a solid boundary, a second vortex ring of opposite sign is formed (*33, 34*). As the two opposite sign vortex rings are in close proximity to each other, they interact and a high pressure region is formed in the area where flows converge and another is formed where an accelerated jet of fluid (VIA) exits from between the two vortices (*31, 33*). Since vortices have below-ambient pressure at their cores, this results in the formation of a 'cloverleaf' of alternating positive and negative pressure fields in close proximity to one another (*33*). Should a propulsive structure be placed at the right orientation within this field significant thrust could be generated. This appears to be what scyphozoan medusae are doing through the formation and positioning of vortices near the propulsive structure (i.e. the bell margin).

The convergence of fluid at the vortex interface results in significantly stronger positive pressure fields along the subumbrellar surface of the bell margin (Figure 3). Negative pressures were greater but not significantly different in cases where a stopping vortex was present

(Supplemental Table 1). Peak negative pressure values always occurred on the aboral side of the bell margin where the starting vortex was forming. The link between greater vorticity/negative pressures and the presence of a stopping vortex are not as direct as the elevated positive pressure observations but may be due to the interaction of the starting and stopping vortices during bell contraction. The highest fluid velocities during the bell contraction occur in a jet formed at the interface of the stopping and starting vortices (Figure 2). In contrast, when no stopping vortex is present the highest fluid velocities occur at on the dorsal side of the bell margin.

While it has been previously recognized that the stopping vortex plays a role in rowing-based medusan propulsion (*4, 19-21*), the mechanistic underpinnings of the vortex-vortex interactions and thrust generation are not well understood. This is like due to the concealed nature of the stopping vortex as it forms and remains either fully or partially obscured under the medusa bell until it interacts, then mostly dissipates and the remnants are ejected in the wake with the starting vortex (*23*). The fact that *A. aurita* exhibits such drastic performance differences in the presence or absence of a stopping vortex—despite starting each swimming cycle from a near-zero velocity and displaying the same kinematics— is evidence of how important stopping vortices are to medusan propulsion. Previous work has demonstrated that the stopping vortex can create additional thrust at the end of a medusa swimming contraction cycle and this contributes to the very low cost of transport observed in this group of animals (*3, 35*). However, the role of the stopping vortex in the subsequent contraction cycle had not been previously considered. Our analysis provides a mechanistic explanation for the enhanced swimming performance (Fig 1) observed in the presence of a well-developed stopping vortex.

**Supplemental Tables:**

Tables

Table 1. Mean values for kinematic and performance metrics for the jellyfish *Aurelia aurita* in the presence of absence of a stopping vortex underneath the animal.

| Parameter | Stopping vortex absent | n | Stopping vortex present | n | P value |
|---|---|---|---|---|---|
| Bell margin speed | 56.3 (s.d. 4.2) mm $s^{-1}$ | 6 | 58.7 (s.d. 4.4) mm $s^{-1}$ | 8 | 0.439 |

| | | | | | |
|---|---|---|---|---|---|
| Bell fineness at peak contraction | 0.549 (s.d. 0.05) | 6 | 0.562 (s.d. 0.04) | 8 | 0.677 |
| Contraction duration | 441.7 (s.d. 7.5) ms | 6 | 428.3 (s.d. 17.2) s | 8 | 0.179 |
| Total displacement | 7.9 (s.d. 1.9) mm | 5 | 12.7 (s.d. 1.5) mm | 7 | *0.007 |
| Maximum swimming speed | 18.2 (s.d. 1.5) mm $s^{-1}$ | 6 | 25.7 (s.d. 0.9) mm $s^{-1}$ | 8 | *<0.001 |
| Maximum negative pressure | -0.34 (s.d. -0.11) Pa | 5 | -0.51 (s.d. 0.12) Pa | 6 | 0.120 |
| Maximum positive pressure | 0.32 (s.d. 0.09) Pa | 5 | 0.74 (s.d. 0.08) Pa | 6 | *<0.001 |
